# DIVAS: an R package for identifying shared and individual variations of multiomics data

**DOI:** 10.64898/2026.01.12.698985

**Authors:** Yinuo Sun, J. S. Marron, Kim-Anh Lê Cao, Jiadong Mao

## Abstract

**Motivation:** Multiomics data integration aims to identify biological patterns shared across molecular modalities. Most existing methods detect either jointly shared variation, across all modalities, or individual variation, unique to a single modality, but overlook partially shared variation, shared by only a subset of modalities. This is a critical limitation, because many biological mechanisms manifest in some but not all molecular modalities.

**Results:** We present an open-source R package implementing DIVAS, a framework for systematically identifying jointly shared, partially shared and individual variations across multiple data types. DIVAS combines angle-based subspace analysis with inference through rotational bootstrap, hierarchically searching all combinations of modalities to decompose multiomics data into interpretable components with scores and loadings. In simulations with a known sharing structure, DIVAS recovered every component across a wide range of noise levels, whereas AJIVE and MOFA+ did not. Applied to multi-modal COVID-19 data, it reveals partially shared immune and metabolic dysregulation patterns underpinning disease severity that conventional approaches would miss.

**Availability and implementation:** DIVAS is available as an R package on GitHub (https://github.com/ByronSyun/DIVAS), with documentation and vignettes at https://byronsyun.github.io/DIVAS/. A step-by-step vignette for the COVID-19 case study is available at https://byronsyun.github.io/DIVAS_COVID19_CaseStudy/.

## 1 Introduction

Modern high-throughput technologies measure diverse molecular layers, from genomics to metabolomics, on the same biological samples. These multiomics datasets offer complementary views of biological processes, yet analysing them in a unified framework remains challenging. The key question is how to systematically identify which biological signals are shared across all omics types (*jointly shared* ), which are shared by only some (*partially shared* ) and which are unique to a single layer (*individual* ).

Most existing multivariate approaches identify only jointly shared variation. Methods based on Canonical Correlation Analysis (CCA) or Partial Least Squares (PLS) find lower-dimensional representations by maximising correlations between input data blocks; widely used examples include Regularised CCA (RCCA) [1], sparse RCCA [2], Regularised Generalised CCA (RGCCA) [3] and DIABLO [4].

Joint and Individual Variation Explained (JIVE) [5] decomposes multiomics data into joint variation shared across all data blocks and individual variation unique to each. Angle-based JIVE (AJIVE) [6] improved on JIVE using principal angle analysis, giving faster joint structure selection and stronger theoretical justification through matrix perturbation theory. Multi-Omics Factor Analysis (MOFA) [7] and MOFA+ [8] use Bayesian group factor analysis to identify latent factors explaining variation within and across omics layers, and began to explore partially shared variation. However, they do not systematically enumerate all possible patterns of partial sharing, and the interpretation of which subset of blocks shares each factor can be ambiguous.

A parallel line of work uses deep generative models, such as MultiVI [9] and MOVE [10]. These target nonlinear representation learning, imputation, batch correction and prediction, returning a learned latent embedding rather than a decomposition labelled by which modalities share each signal; they are therefore complementary to the methods considered here.

Recent methods target partially shared variation directly. Structural Learning and Integrative Decomposition (SLIDE) [11] models variation shared between specific subsets of blocks using structured sparsity, with block-specific patterns determined by bi-cross-validation, but it does not systematically detect all patterns of partial sharing and lacks statistical inference for those it finds. Hierarchical Nuclear Norm (HNN) penalisation [12] formulates partially shared variation through hierarchical levels, using a convex formulation that avoids scores-loadings factorisation; it offers identifiability guarantees but requires computationally intensive tuning parameter selection. No package is available for either method.

Table 1 summarises these methods. The critical gap is the systematic identification of partially shared variation, which matters most with three or more modalities: there is no guarantee that the most interesting biological patterns are captured by jointly shared variation alone. A regulatory mechanism may affect transcriptomics and proteomics but not metabolomics, while metabolic dysregulation may appear in metabolomics and proteomics without being reflected transcriptionally.

**Table 1.** Comparison of multiomics integration methods. ‘✓’ indicates that a method is capable of performing a specific task, while ‘✗’ denotes that the method is not equipped to handle the task. The capabilities of DIVAS, MOFA+ and AJIVE are additionally assessed empirically in Supplementary Methods Section S3.

| Method | Jointly shared | Partially shared | Individual | Statistical inference | Package availability |
| --- | --- | --- | --- | --- | --- |
| RCCA/RGCCA | ✓ | ✗ | ✗ | ✗ | CRAN [15] |
| DIABLO | ✓ | ✗ | Limited<sup>a</sup> | ✗ | Bioconductor [16] |
| JIVE | ✓ | ✗ | ✓ | Limited<sup>b</sup> | CRAN [17] |
| AJIVE | ✓ | ✗ | ✓ | Limited<sup>c</sup> | GitHub [18] |
| MOFA/MOFA+ | ✓ | Limited<sup>d</sup> | ✓ | ✓<sup>e</sup> | Bioconductor [19] |
| SLIDE | ✓ | ✓ | ✓ | Limited<sup>f</sup> | N/A |
| HNN | ✓ | ✓ | ✓ | Limited<sup>g</sup> | N/A |
| **DIVAS** | ✓ | ✓ | ✓ | ✓ | GitHub [20] |

<sup>a</sup> Focus on joint variation; individual variation not explicitly modelled
<sup>b</sup> Permutation testing for rank selection only
<sup>c</sup> Angle bounds for joint vs individual separation, but not for rank selection
<sup>d</sup> Learns sparsity patterns but does not detect all partial sharing patterns; the interpretation of shared patterns can be ambiguous
<sup>e</sup> Bayesian inference on factors and loadings
<sup>f</sup> Cross-validation for model selection only
<sup>g</sup> Convex optimization with tuning parameter selection via bi-cross-validation

Addressing this gap, Data Integration Via Analysis of Subspaces (DIVAS) was proposed by Prothero et al. [13] (see also Ackerman et al. [14] for an application in neuroimaging). The framework systematically identifies jointly shared, partially shared and individual variations, provides inference procedures with statistical guarantees, and uses broadly useful defaults enabling fully data-driven integration without manual tuning.

Until now, however, DIVAS has been available only as closed-source MATLAB code, limiting its uptake. The statistical methodology is that of Prothero et al. and is not modified here; the contribution of the present work is software: (i) an open-source R implementation of the full algorithm; (ii) visualisation and downstream-interpretation tools not previously available; (iii) documentation and vignettes for users without a background in matrix perturbation theory or convex optimisation; and (iv) the first quantitative benchmark of DIVAS against alternative methods on simulated data with a known sharing structure, together with an applied multiomic case study.

## 2 Methods

The input of DIVAS is a collection of *K* data blocks *X*_1_, …, *X*_*K*_ from *N* samples, where each *X*_*k*_ is a (*d*_*k*_ *× N* ) feature-by-sample matrix. These data blocks represent multiomic measurements from the same samples. The DIVAS algorithm, detailed in Prothero et al. [13], comprises the following two main steps:

1. Denoising individual data blocks;
2. Identifying shared and individual structures among denoised data blocks.

Both steps are fully data-driven and tuning free. The output is a set of components explaining shared and individual variations, each with scores and loadings as in standard factor models; see Fig 1 for a schema.

**Fig 1.**
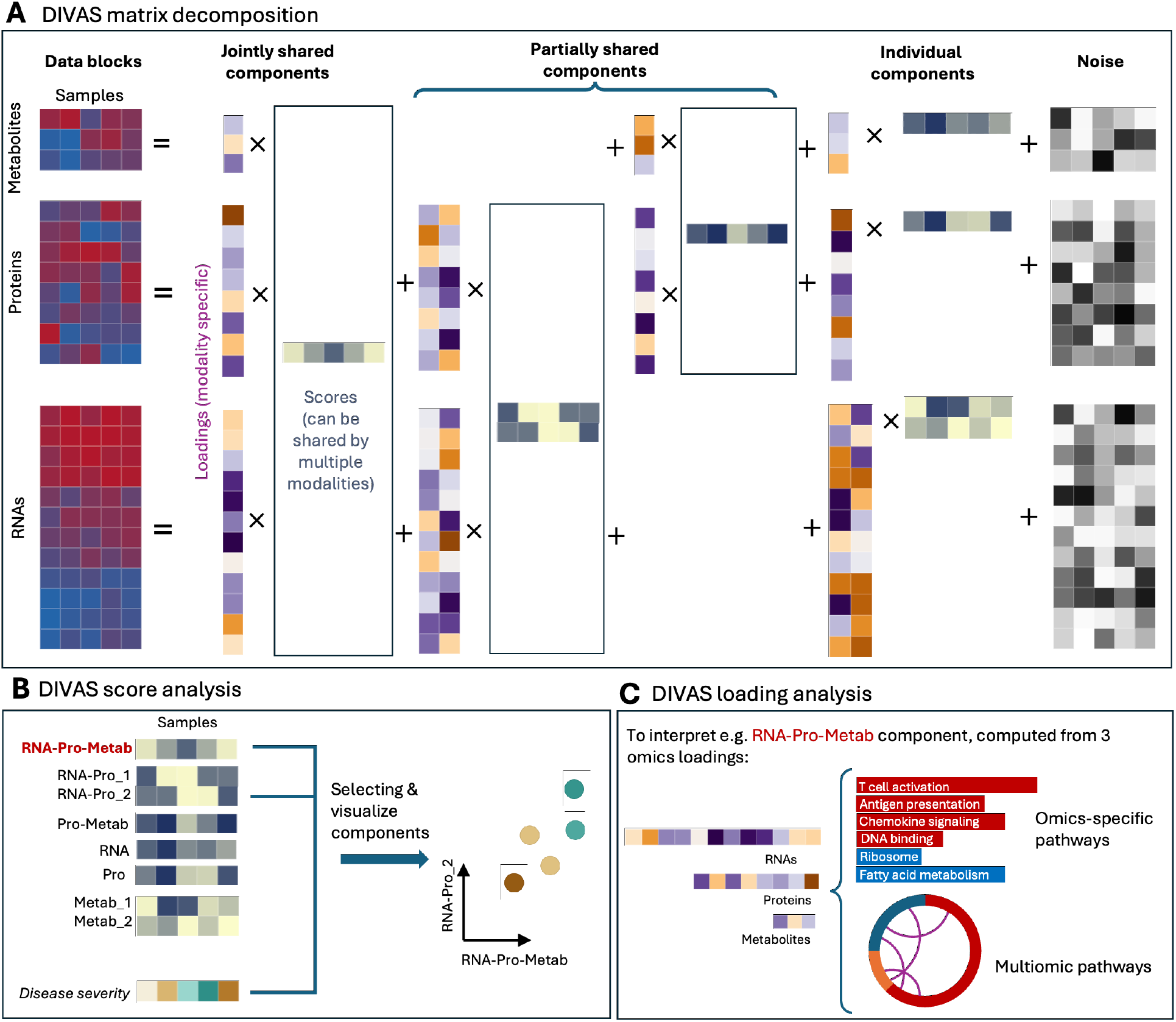
Schema of DIVAS. **A** DIVAS decomposes input data blocks into components representing jointly shared, partially shared and individual modes of variation. Here we present a toy example taking transcriptomics, proteomics and metabolomics as input. Each DIVAS component, representing a specific mode of variation, is defined by a score vector and one or more loading vectors (corresponding to one or more modalities). In the toy example, we found one 3-way jointly shared component (RNA–protein–metabolite), three 2-way partially shared components (two RNA–protein, one protein–metabolite) and four individual components (two RNA, one protein, one metabolite). **B** DIVAS scores are lower dimensional summaries of samples and can be used for visualisation. For example, we can select the scores most highly correlated with disease severity and visualise them using a scatter plot. **C** DIVAS loadings measure the contribution of individual omics features to scores. Whereas one DIVAS component only has one common score vector, it may have multiple loading vectors, corresponding a set of data blocks. By identifying pathways represented by important omics features, we can interpret the biological meanings of individual components. In the notation of Section 2, each data block *X*_*k*_ in **A** is *d*_*k*_ *× N*, each loading matrix *L*_***i***,*k*_ is *d*_*k*_ *× r*_***i***_, each score matrix *S****i***^⊤^ is *r*_***i***_ *× N* and each noise matrix *E*_*k*_ is *d*_*k*_*× N* . The rank decomposition and angle diagnostics are returned directly by `DIVASmain`; the score visualisations in **B** are produced by the companion vizOmics package, which takes DIVAS output objects as input; and for **C**, DIVAS::getTopFeatures returns ranked feature tables that users pass to their preferred enrichment and plotting tools.

### Denoising data blocks

We assume that each data block has an underlying low rank structure *A*_*k*_, which is contaminated by observational noise *E*_*k*_ (see Fig 1A). That is, we assume *X*_*k*_ = *A*_*k*_ + *E*_*k*_. To recover *A*_*k*_ from noisy observation *X*_*k*_, we apply Principal Component Analysis (PCA) to *X*_*k*_ and take the first PCs. The number of PCs to use is determined by an optimal shrinkage method [21, 22] in a fully data-driven way.

### Identifying shared and individual structures

We assume that the total variation in each *A*_*k*_, the noiseless version of the *k*th data block, can be decomposed into jointly shared, partially shared and individual variations. That is, with ***i*** denoting a subset of the index set of all data blocks *{*1, …, *K}*, we assume that

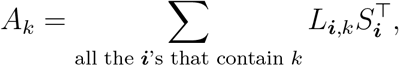

where *S*_***i***_ is an (*N × r*_***i***_) score matrix, with *r*_***i***_ denoting the number of components shared by data blocks in ***i***, and where *L*_***i***,*k*_ is a (*d*_*k*_ *× r*_***i***_) loading matrix, linking the *r*_***i***_ components with the *d*_*k*_ features of the *k*th data block. That is, a DIVAS component shared by data blocks in ***i*** comprises *common* scores for all those data blocks, together with one loading vector for each data block in ***i***; see Fig 1A for an example involving 3 data blocks.

The core of the algorithm is a convex-concave optimisation procedure [13], of which we give only a sketch here. A shared structure represents a pattern of biological variation capturing how omics variables co-vary across samples, and finding it amounts to identifying the directions pointing towards the most important such patterns. DIVAS searches hierarchically, starting from the highest-order structures, shared among the most data blocks, and moving progressively to lower-order and individual structures. For each candidate direction it tests whether that direction stays close to the signal patterns of all included blocks whilst remaining sufficiently distant from the excluded ones, continuing until no further directions can be found for the current structure. The strength of this approach is that it determines automatically how many dimensions are needed for each type of relationship, giving a data-driven decomposition into interpretable shared and individual components.

### Interpreting scores and loadings

The decomposition yields scores and loadings for each identified component. Each component has a score vector summarising, in one dimension, how samples vary along that mode of variation, shared across all data blocks involved in the structure (Fig 1A&B). A jointly shared component across transcriptomics, proteomics and metabolomics, for example, has a single common score vector.

Loadings link those shared scores to the features measured in each block. A component has one score vector but may have several modality-specific loading vectors (Fig 1A&C), each indicating which features of that block contribute to the shared variation. For a jointly shared component, the transcriptomics loadings reveal which genes drive the pattern, while the proteomics and metabolomics loadings identify the associated proteins and metabolites. The top contributing features can then be taken forward for pathway analysis (Fig 1C).

## 3 Results

### Benchmarking on simulated data

We first assessed whether DIVAS recovers a known sharing structure, using three matched data blocks generated with one jointly shared, three partially shared and three individual rank-one components, over ten noise levels and five random seeds (Supplementary Methods Section S3). DIVAS identified the correct rank for all seven components, in every seed, up to a noise standard deviation of 2.00, and the chordal distance to the true score subspaces increased smoothly with noise (Supplementary Figs S3 and S4). On the same data, AJIVE over-estimated the rank of every component it returned and provides no partially shared components at all, while MOFA+ factors, mapped post hoc to sharing subsets by their per-block variance explained, recovered the partially shared components in at most 4 of 50 datasets (Supplementary Figs S5 and S6).

### Study design and data

We analysed a multiomic dataset from a cohort of mild to severe COVID-19 patients [39]. The study included 139 patients measured at two time points, T1 and T2, giving 265 samples. Of the available omics types we selected scRNA-seq, bulk proteomics and bulk metabolomics; 120 patients had all three, giving 240 samples across the two time points. Since DIVAS assumes independence between samples for inference, we avoided repeated measurements of the same subjects. We used stratified sampling to split the 120 patients into two groups of 60, retaining the T1 observation of one group and the T2 observation of the other, so that each patient contributes exactly one sample. After restricting to samples with complete measurements in all six data blocks, 114 samples remained.

DIVAS requires only that its input blocks are measured on the same samples in the same order; it does not require any particular type of block. Here, however, the scRNA-seq measurements are cell-level whereas the proteomics and metabolomics are sample-level. To harmonise this difference in granularity, we annotated cell types with CellTypist [40, 41] and pseudo-bulked CD4 T, CD8 T, CD14 monocyte and natural killer (NK) cells within each sample, these being abundant PBMC populations with robust annotations and important roles in the COVID-19 immune response [39]. The gene expression (GEX) profiles of these 4 cell types, together with proteomics (pro) and metabolomics (metab), formed 6 input blocks. Splitting a modality by cell type is a design choice specific to this case study, not a required preprocessing step: it lets DIVAS resolve programs shared across cellular compartments as well as across molecular modalities. See Supplementary Materials Section S1 for preprocessing and cell type annotation details.

### Biological insights

Applying DIVAS to the 6-modal COVID-19 data, we found over 90 statistically significant components, ranging from 6-way (jointly shared variation) down to 1-way (individual variation) (see Supplementary Fig S1). We then ranked all DIVAS component according to the correlation between their scores with the COVID-19 severity score, an ordinal variable ranging from 1 to 7. The top two most highly correlated components are visualised in Fig 2A (see Supplementary Fig S2 for the top five components).

**Fig 2.**
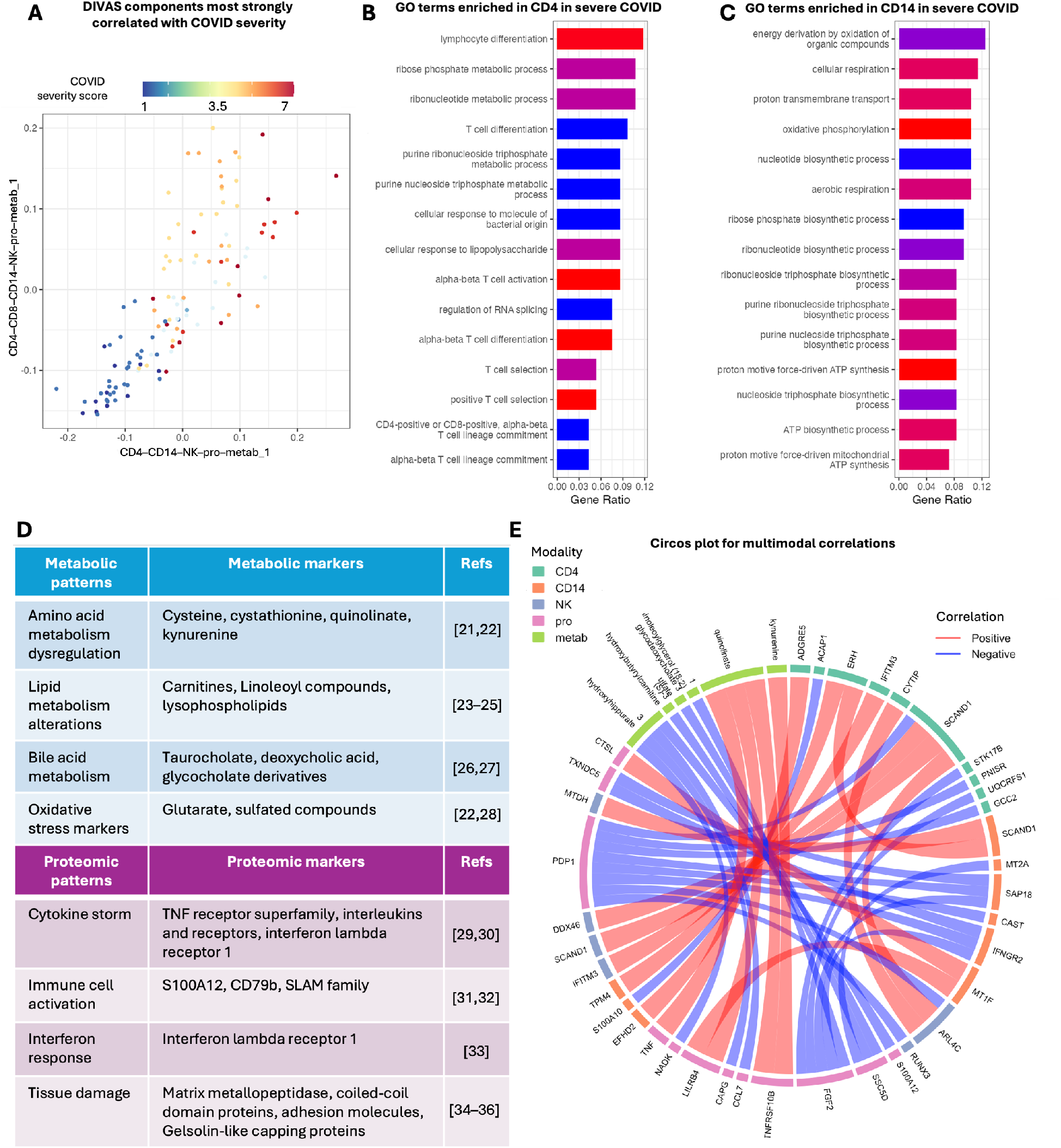
Uncovering multiomic pathways underpinning COVID-19 severity. **A** Visualising the scores of DIVAS components most strongly correlated with COVID severity. A component, e.g. CD4– CD14–NK–pro–metab 1, is named after the data blocks it represents. As there may be multiple components corresponding to a particular combination of data blocks, we also use an index to distinguish them. **B&C** Gene ontology (GO) terms enriched in CD4 T cell and CD14 monocyte gene expressions in severe COVID-19. **D** Metabolic and proteomic patterns observed in severe COVID-19 [23–38]. **E** Circos plot illustrating within- and cross-modal correlations, where the thickness of ribbon represents the absolute value of pairwise Spearman correlation coefficient between connected features.

DIVAS loadings rank modality-specific features by their contribution to a component of interest. For CD4–CD14–NK–pro–metab 1, the component most strongly correlated with severity, we took the CD4 GEX features most positively and negatively associated with severity and conducted gene set overrepresentation analysis (Supplementary Methods Section S2). The results showed marked differences in CD4 T cell phenotype between severe and mild COVID-19: mild patients showed a more coordinated immune response, marked by T cell and hematopoiesis regulations (Fig 2B), whereas severe patients showed a highly activated but dysfunctional CD4 T cell state, marked by metabolic stress (Fig 2C). From the proteomics and metabolomics modalities, we found markers representing a series of immune and metabolic dysregulations (Fig 2D), confirming the CD4 T cell dysregulation seen in Figs 2B&C from complementary omics perspectives. The pathways in Figs 2B– D are well documented in severe COVID-19; recovering them is a positive control that the decomposition is biologically sound. What DIVAS reveals is how these pathways are shared: these programs emerged not as a jointly shared signal across all six blocks, but as a single partially shared component spanning CD4 T, CD14 monocyte, NK, proteomic and metabolomic blocks while excluding CD8 T cells. A method restricted to jointly shared variation would have to attribute this signal to all six blocks or to none.

Finally, we identified the top features from all 5 modalities involved in the CD4–CD14– NK–pro–metab 1 component and examined their correlations (see Supplementary Methods Section S2). Interestingly, we identified correlation structures that helped explain the immune dysregulation mentioned above. Namely multiple chemokines (e.g. CCL7), TNF receptors (e.g. TNFRSF10B) and metabolites (e.g. quinolinate) formed an interconnected network (Fig 2E), suggesting the cytokine storm that is commonly observed in severe COVID-19 immune response [31, 32, 35]. Another highlight was the multiple strong negative correlations identified across modalities (blue links in Fig 2E). For example, PDP1 protein was found to be negatively correlated with IFNGR2 gene in CD14 monocytes. This suggested a discordance in severe COVID-19 patients between myeloid activation, for which IFNGR2 is essential [42], and energy production via oxidative metabolism, promoted by PDP1 [43].

## 4 Discussion

We introduced the DIVAS R package, making available to the bioinformatics community a framework that systematically identifies jointly shared, partially shared and individual variations across multiple data blocks. This fills a critical gap: many biological mechanisms are reflected in only some molecular modalities, and methods restricted to fully shared or individual variation miss these intermediate patterns. Our benchmark confirms this concretely, with neither AJIVE nor MOFA+ reliably recovering the partially shared components that DIVAS identified correctly across almost the entire noise range tested.

The properties underpinning this performance are: (1) hierarchical search through all combinations of modalities using angle-based subspace analysis; (2) built-in inference through principal angle analysis and rotational bootstrap; (3) simultaneous consideration of sample and feature spaces; and (4) handling of high-dimensional data through optimal shrinkage.

DIVAS is complementary to probabilistic latent factor models such as MOFA+ [8] and deep generative frameworks such as MultiVI [9] and MOVE [10]. These are preferable when the goal is large-scale nonlinear representation learning, imputation, batch correction or prediction. DIVAS is preferable when the question is which modalities share a signal, and the answer must be explicit, interpretable and accompanied by subset-specific ranks, scores, loadings and inference. The two can be combined: DIVAS loadings offer a structure-aware way to prioritise features from biologically relevant components before predictive modelling, in place of generic filters such as variance thresholding or univariate selection.

Two practical considerations should guide use. First, stable decomposition depends not only on sample size but also on noise, feature filtering, preprocessing quality and the stability of the initial rank estimation, so we do not recommend a universal minimum sample size. Our simulations and case study indicate that DIVAS suits matched studies with roughly 100–300 samples and 3–6 blocks, with stability checked by repeated runs and perturbation diagnostics near those limits. Notably, DIVAS also yielded interpretable results in preliminary analyses of a four-omics dataset comprising 19 samples, although these analyses are not presented here. Runtime of DIVAS grows with the number of blocks, as the candidate subsets grow combinatorially. Second, normalisation and centring affect each block’s relative contribution and hence the estimated subspaces: `DIVASmain` exposes `colCent` and `rowCent`, both `FALSE` by default, with row-centring appropriate when feature-level mean differences should be removed beforehand. Since DIVAS assumes blocks measured on the same samples in the same order, reproducibility also depends on careful block construction upstream; our vignettes document this workflow end to end, including the diagnostics to inspect before interpreting components.

The package is designed for researchers without specialised expertise in matrix perturbation theory or convex optimisation, providing straightforward functions for decomposition and downstream interpretation. As multiomics studies grow more common and more complex, DIVAS offers a principled way to extract the full spectrum of biological information encoded across complementary molecular measurements.

## Author contributions

JM and JSM conceived the project; JM and KALC secured funding; YS developed the R package and tested it on simulated and real-world datasets under the supervision of JM; JM and YS performed data analyses for the COVID-19 case study and compiled the vignette; JM and YS wrote the manuscript; all authors participated in revising the manuscript.

## Data availability

The raw scRNA-seq count tables for the COVID-19 case study can be accessed by Array Express under the accession number E-MTAB-9357; metabolomic and proteomic datasets are available from Mendeley Data [44]. Processed data used in this paper are available from Zenodo [45].

## Code availability

The DIVAS R package is available on GitHub [20], with function reference documentation and vignettes at https://byronsyun.github.io/DIVAS/. The pipeline used for processing COVID-19 data is available on GitHub [46]. The code for reproducing the results and simulation benchmarks in this paper is available on GitHub [47].

## Acknowledgements

We would like to thank Dr Saritha Kodikara (The University of Melbourne) and Dr Geraldine Kong (Peter Doherty Institute) for helpful discussions.

## Use of large language model (LLM)

For generating the summary of metabolic and proteomic pathways in Fig 2D, Claude Sonnet 4.5 was used to identify potential pathways based on features with top DIVAS loading values. These pathways were then validated by a thorough literature search and review. We also used Claude Opus 4.8 and ChatGPT 5.6 Sol to check grammatical mistakes and textual inconsistencies.

## References

[1] Arthur Tenenhaus and Michel Tenenhaus. Regularized generalized canonical correlation analysis. Psychometrika, 76(2):257–284, 2011.

[2] Arthur Tenenhaus, Cathy Philippe, Vincent Guillemot, Kim-Anh Lê Cao, Jacques Grill, and Vincent Frouin. Variable selection for generalized canonical correlation analysis. Biostatistics, 15(3):569–583, 2014.

[3] Michel Tenenhaus, Arthur Tenenhaus, and Patrick J F Groenen. Regularized generalized canonical correlation analysis: A framework for sequential multiblock component methods. Psychometrika, 82(3):737–777, 2017.

[4] Amrit Singh, Casey P Shannon, Benoît Gautier, Florian Rohart, Michäel Vacher, Scott J Tebbutt, and Kim-Anh Lê Cao. DIABLO: an integrative approach for identifying key molecular drivers from multi-omics assays. Bioinformatics, 35(17):3055–3062, 2019.

[5] Eric F Lock, Katherine A Hoadley, J S Marron, and Andrew B Nobel. Joint and individual variation explained (JIVE) for integrated analysis of multiple data types. Ann. Appl. Stat., 7 (1):523–542, 2013.

[6] Qing Feng, Meilei Jiang, Jan Hannig, and J S Marron. Angle-based joint and individual variation explained. J. Multivar. Anal., 166:241–265, 2018.

[7] Ricard Argelaguet, Britta Velten, Damien Arnol, Sascha Dietrich, Thorsten Zenz, John C Marioni, Florian Buettner, Wolfgang Huber, and Oliver Stegle. Multiomics factor analysis-a framework for unsupervised integration of multi-omics data sets. Mol. Syst. Biol., 14(6):e8124, 2018.

[8] Ricard Argelaguet, Damien Arnol, Danila Bredikhin, Yonatan Deloro, Britta Velten, John C Marioni, and Oliver Stegle. MOFA+: a statistical framework for comprehensive integration of multi-modal single-cell data. Genome Biol., 21(1):111, 2020.

[9] Tal Ashuach, Mariano I. Gabitto, Rohan V. Koodli, Giuseppe-Antonio Saldi, Michael I. Jordan, and Nir Yosef. MultiVI: deep generative model for the integration of multimodal data. Nature Methods, 20(8):1222–1231, 2023. URL 10.1038/s41592-023-01909-9.

[10] Rosa Lundbye Allesøe, Agnete Troen Lundgaard, Ricardo Hernández Medina, Alejandro Aguayo-Orozco, Joachim Johansen, Jakob Nybo Nissen, Caroline Brorsson, Gianluca Mazzoni, Lili Niu, Jorge Hernansanz Biel, Cristina Leal Rodríguez, Valentas Brasas, Henry Webel, Michael Eriksen Benros, Anders Gorm Pedersen, Piotr Jaroslaw Chmura, Ulrik Plesner Jacobsen, Andrea Mari, Robert Koivula, Anubha Mahajan, Ana Viñuela, Juan Fernandez Tajes, Sapna Sharma, Mark Haid, Mun-Gwan Hong, Petra B. Musholt, Federico De Masi, Josef Vogt, Helle Krogh Pedersen, Valborg Gudmundsdottir, Angus Jones, Gwen Kennedy, Jimmy Bell, E. Louise Thomas, Gary Frost, Henrik Thomsen, Elizaveta Hansen, Tue Haldor Hansen, Henrik Vestergaard, Mirthe Muilwijk, Marieke T. Blom, Leen M. ‘t Hart, Francois Pattou, Violeta Raverdy, Soren Brage, Tarja Kokkola, Alison Heggie, Donna McEvoy, Miranda Mourby, Jane Kaye, Andrew Hattersley, Timothy McDonald, Martin Ridderstråle, Mark Walker, Ian Forgie, Giuseppe N. Giordano, Imre Pavo, Hartmut Ruetten, Oluf Pedersen, Torben Hansen, Emmanouil Dermitzakis, Paul W. Franks, Jochen M. Schwenk, Jerzy Adamski, Mark I. McCarthy, Ewan Pearson, Karina Banasik, Simon Rasmussen, Søren Brunak, and IMI DIRECT Consortium. Discovery of drug-omics associations in type 2 diabetes with generative deep-learning models. Nature Biotechnology, 41(3):399–408, 2023. URL 10.1038/s41587-022-01520-x.

[11] Irina Gaynanova and Gen Li. Structural learning and integrative decomposition of multi-view data. Biometrics, 75(4):1121–1132, 2019.

[12] Sangyoon Yi, Raymond Ka Wai Wong, and Irina Gaynanova. Hierarchical nuclear norm penalization for multi-view data integration. Biometrics, 79(4):2933–2946, 2023.

[13] Jack Prothero, Meilei Jiang, Jan Hannig, Quoc Tran-Dinh, Andrew Ackerman, and J S Marron. Data integration via analysis of subspaces (DIVAS). TEST, 33:633–674, 2024. URL 10.1007/s11749-024-00923-z.

[14] Andrew Ackerman, Zhengwu Zhang, Jan Hannig, Jack Prothero, and JS Marron. Multifaceted neuroimaging data integration via analysis of subspaces. Psychometrika, pages 1–26, 2024.

[15] Fabien Girka, Etienne Camenen, Caroline Peltier, Arnaud Gloaguen, Vincent Guillemot, Laurent Le Brusquet, and Arthur Tenenhaus. RGCCA: Regularized and Sparse Generalized Canonical Correlation Analysis for Multiblock Data, 2023. URL https://CRAN.R-project.org/package=RGCCA. R package version 3.0.3.

[16] Kim-Anh Lê Cao, Florian Rohart, Ignacio Gonzalez, Sebastien Dejean, Al J Abadi, Max Bladen, Benoit Gautier, Francois Bartolo, Pierre Monget, Jeff Coquery, FangZou Yao, Benoit Liquet, and Eva Hamrud. mixOmics: Omics Data Integration Project, 2025. URL https://www.bioconductor.org/packages/release/bioc/html/mixOmics.html. Bioconductor Release 3.22.

[17] Michael J. O’Connell, Eric F. Lock, and Adam Kaplan. r.jive: Perform JIVE Decomposition for Multi-Source Data, 2020. URL https://CRAN.R-project.org/package=r.jive. R package version 2.4.

[18] Iain Carmichael. idc9/r.jive: First GitHub release, 2020. URL 10.5281/zenodo.4091755. R package version 0.0.1.

[19] Ricard Argelaguet, Damien Arnol, Danila Bredikhin, and Britta Velten. MOFA2: Multi-Omics Factor Analysis v2, 2025. URL https://www.bioconductor.org/packages/release/bioc/html/MOFA2.html. Bioconductor Release 3.22.

[20] Yinuo Sun and Jiadong Mao. DIVAS: Data Integration via Analysis of Subspaces, 2025. URL https://github.com/ByronSyun/DIVAS. R package version 0.1.1, GitHub repository.

[21] Matan Gavish and David L Donoho. Optimal shrinkage of singular values. IEEE Trans. Inf. Theory, 63(4):2137–2152, 2017. doi: 10.1109/TIT.2017.2653801.

[22] Andrey A Shabalin and Andrew B Nobel. Reconstruction of a low-rank matrix in the presence of gaussian noise. J. Multivar. Anal., 118:67–76, 2013. URL 10.1016/j.jmva.2013.03.005.

[23] Murat Cihan, Ö zlem Doğan, Ceyhan Ceran Serdar, Arzu Altunçekiç Yildirim Celali Kurt, and Muhittin A Serdar. Kynurenine pathway in coronavirus disease (COVID-19): Potential role in prognosis. J. Clin. Lab. Anal., 36(3):e24257, 2022.

[24] Tiffany Thomas, Davide Stefanoni, Julie A Reisz, Travis Nemkov, Lorenzo Bertolone, Richard O Francis, Krystalyn E Hudson, James C Zimring, Kirk C Hansen, Eldad A Hod, Steven L Spitalnik, and Angelo D’Alessandro. COVID-19 infection alters kynurenine and fatty acid metabolism, correlating with IL-6 levels and renal status. JCI Insight, 5(14), 2020.

[25] Vamsi P Guntur, Travis Nemkov, Esther de Boer, Michael P Mohning, David Baraghoshi, Francesca I Cendali, Inigo San-Millán, Irina Petrache, and Angelo D’Alessandro. Signatures of mitochondrial dysfunction and impaired fatty acid metabolism in plasma of patients with post-acute sequelae of COVID-19 (PASC). Metabolites, 12(11):1026, 2022.

[26] Chunyu Li, Ruwei Ou, Qianqian Wei, and Huifang Shang. Carnitine and COVID-19 susceptibility and severity: A mendelian randomization study. Front. Nutr., 8:780205, 2021.

[27] Juntong Wei, Xiaoyu Liu, Weimin Xiao, Jiahua Lu, Li Guan, Zhangfu Fang, Jiaping Chen, Baoqing Sun, Zongwei Cai, Xizhuo Sun, Hua-Ling Chen, Nanshan Zhong, Zhigang Liu, Jun Yang, Xiaojun Xiao, and Shau-Ku Huang. Phospholipid remodeling and its derivatives are associated with COVID-19 severity. J. Allergy Clin. Immunol., 151(5):1259–1268, 2023.

[28] Felipe N Piñol Jiménez, Virginia Capó-de Paz, Julián F Ruiz-Torres, Teresita Montero-González, Sara A Urgellés-Carreras, Andrés Breto-García, Armando Amador-Armenteros, María M Llerena-Mesa, and Ana G Galarraga-Lazcano. High levels of serum bile acids in COVID-19 patients on hospital admission. MEDICC Rev., 24(3–4):53–56, 2022.

[29] Xiaoru Huang, Xuening Liu, and Zijian Li. Bile acids and coronavirus disease 2019. Acta Pharm. Sin. B., 14(5):1939–1950, 2024.

[30] Carlos A Labarrere and Ghassan S Kassab. Glutathione deficiency in the pathogenesis of SARS-CoV-2 infection and its effects upon the host immune response in severe COVID-19 disease. Front. Microbiol., 13:979719, 2022.

[31] Christian Zanza, Tatsiana Romenskaya, Alice Chiara Manetti, Francesco Franceschi, Raffaele La Russa, Giuseppe Bertozzi, Aniello Maiese, Gabriele Savioli, Gianpietro Volonnino, and Yaroslava Longhitano. Cytokine storm in COVID-19: Immunopathogenesis and therapy. Medicina (Kaunas), 58(2):144, 2022.

[32] Shintaro Hojyo, Mona Uchida, Kumiko Tanaka, Rie Hasebe, Yuki Tanaka, Masaaki Murakami, and Toshio Hirano. How COVID-19 induces cytokine storm with high mortality. Inflamm. Regen., 40(1):37, 2020.

[33] Patricia Mester, Dennis Keller, Claudia Kunst, Ulrich Räth, Sophia Rusch, Stephan Schmid, Sabrina Krautbauer, Martina Müller, Christa Buechler, and Vlad Pavel. High serum S100A12 as a diagnostic and prognostic biomarker for severity, multidrug-resistant bacteria superinfection and herpes simplex virus reactivation in COVID-19. Viruses, 16(7):1084, 2024.

[34] Cristiana Iosef, Claudio M Martin, Marat Slessarev, Carolina Gillio-Meina, Gediminas Cepinskas, Victor K M Han, and Douglas D Fraser. COVID-19 plasma proteome reveals novel temporal and cell-specific signatures for disease severity and high-precision disease management. J. Cell. Mol. Med., 27(1):141–157, 2023.

[35] Santhamani Ramasamy and Selvakumar Subbian. Critical determinants of cytokine storm and type I interferon response in COVID-19 pathogenesis. Clin. Microbiol. Rev., 34(3), 2021.

[36] Monica Gelzo, Sara Cacciapuoti, Biagio Pinchera, Annunziata De Rosa, Gustavo Cernera, Filippo Scialò, Marika Comegna, Mauro Mormile, Gabriella Fabbrocini, Roberto Parrella, Gaetano Corso, Ivan Gentile, and Giuseppe Castaldo. Matrix metalloproteinases (MMP) 3 and 9 as biomarkers of severity in COVID-19 patients. Sci. Rep., 12(1):1212, 2022.

[37] Rebecca Salomão, Victoria Assis, Ivo Vieira de Sousa Neto, Bernardo Petriz, Nicolas Babault, João Luiz Quaglioti Durigan, and Rita de Cássia Marqueti. Involvement of matrix metalloproteinases in COVID-19: Molecular targets, mechanisms, and insights for therapeutic interventions. Biology (Basel), 12(6):843, 2023.

[38] Cody B Jackson, Michael Farzan, Bing Chen, and Hyeryun Choe. Mechanisms of SARS-CoV-2 entry into cells. Nat. Rev. Mol. Cell Biol., 23(1):3–20, 2022.

[39] Yapeng Su, Daniel Chen, Dan Yuan, Christopher Lausted, Jongchan Choi, Chengzhen L Dai, Valentin Voillet, Venkata R Duvvuri, Kelsey Scherler, Pamela Troisch, Priyanka Baloni, Guangrong Qin, Brett Smith, Sergey A Kornilov, Clifford Rostomily, Alex Xu, Jing Li, Shen Dong, Alissa Rothchild, Jing Zhou, Kim Murray, Rick Edmark, Sunga Hong, John E Heath, John Earls, Rongyu Zhang, Jingyi Xie, Sarah Li, Ryan Roper, Lesley Jones, Yong Zhou, Lee Rowen, Rachel Liu, Sean Mackay, D Shane O’Mahony, Christopher R Dale, Julie A Wallick, Heather A Algren, Michael A Zager, ISB-Swedish COVID19 Biobanking Unit, Wei Wei, Nathan D Price, Sui Huang, Naeha Subramanian, Kai Wang, Andrew T Magis, Jenn J Hadlock, Leroy Hood, Alan Aderem, Jeffrey A Bluestone, Lewis L Lanier, Philip D Greenberg, Raphael Gottardo, Mark M Davis, Jason D Goldman, and James R Heath. Multiomics resolves a sharp disease-state shift between mild and moderate COVID-19. Cell, 183(6):1479–1495.e20, 2020. URL https://www.cell.com/cell/fulltext/S0092-8674(20)31444-6?_returnUR=.

[40] C Domínguez Conde, C Xu, L B Jarvis, D B Rainbow, S B Wells, T Gomes, S K Howlett, O Suchanek, K Polanski, H W King, L Mamanova, N Huang, P A Szabo, L Richardson, L Bolt, E S Fasouli, K T Mahbubani, M Prete, L Tuck, N Richoz, Z K Tuong, L Campos, H S Mousa, E J Needham, S Pritchard, T Li, R Elmentaite, J Park, E Rahmani, D Chen, D K Menon, O A Bayraktar, L K James, K B Meyer, N Yosef, M R Clatworthy, P A Sims, D L Farber, K Saeb-Parsy, L L Jones, and S A Teichmann. Cross-tissue immune cell analysis reveals tissue-specific features in humans. Science, 376(6594):eabl5197, 2022.

[41] Chuan Xu, Martin Prete, Simone Webb, Laura Jardine, Benjamin J Stewart, Regina Hoo, Peng He, Kerstin B Meyer, and Sarah A Teichmann. Automatic cell-type harmonization and integration across human cell atlas datasets. Cell, 186(26):5876–5891.e20, 2023.

[42] Nataša Todorović-Raković and Jonathan R Whitfield. Between immunomodulation and immunotolerance: The role of IFNγ in SARS-CoV-2 disease. Cytokine, 146(155637):155637, 2021.

[43] Vikalp Kumar and Miriam L Greenberg. Emerging roles of pyruvate dehydrogenase phosphatase 1: a key player in metabolic health. Front. Physiol., 16:1596636, 2025.

[44] Yapeng Su. Multi-omics resolves a sharp disease-state shift between mild and moderate COVID-19. Su et al, 2020. URL https://data.mendeley.com/datasets/tzydswhhb5/5. Mendeley Data, V5.

[45] Yinuo Sun. COVID-19 Multi-omics Data Integration using DIVAS: scRNA-seq, Proteomics, and Metabolomics from 114 Patient Samples, 2025. URL 10.5281/zenodo.17430294. Zenodo Dataset.

[46] Yinuo Sun. DIVAS COVID-19 Case Study, 2025. URL https://github.com/ByronSyun/DIVAS_COVID19_CaseStudy. GitHub Repository.

[47] Jiadong Mao and Yinuo Sun. Finding multiomic markers and pathways underpinning COVID-19 severity using DIVAS, 2025. URL https://byronsyun.github.io/DIVAS_COVID19_CaseStudy/. R Markdown Vignette.

