## Supplemental Figures for "DIVAS: an R package for identifying shared and individual variations of multiomics data"

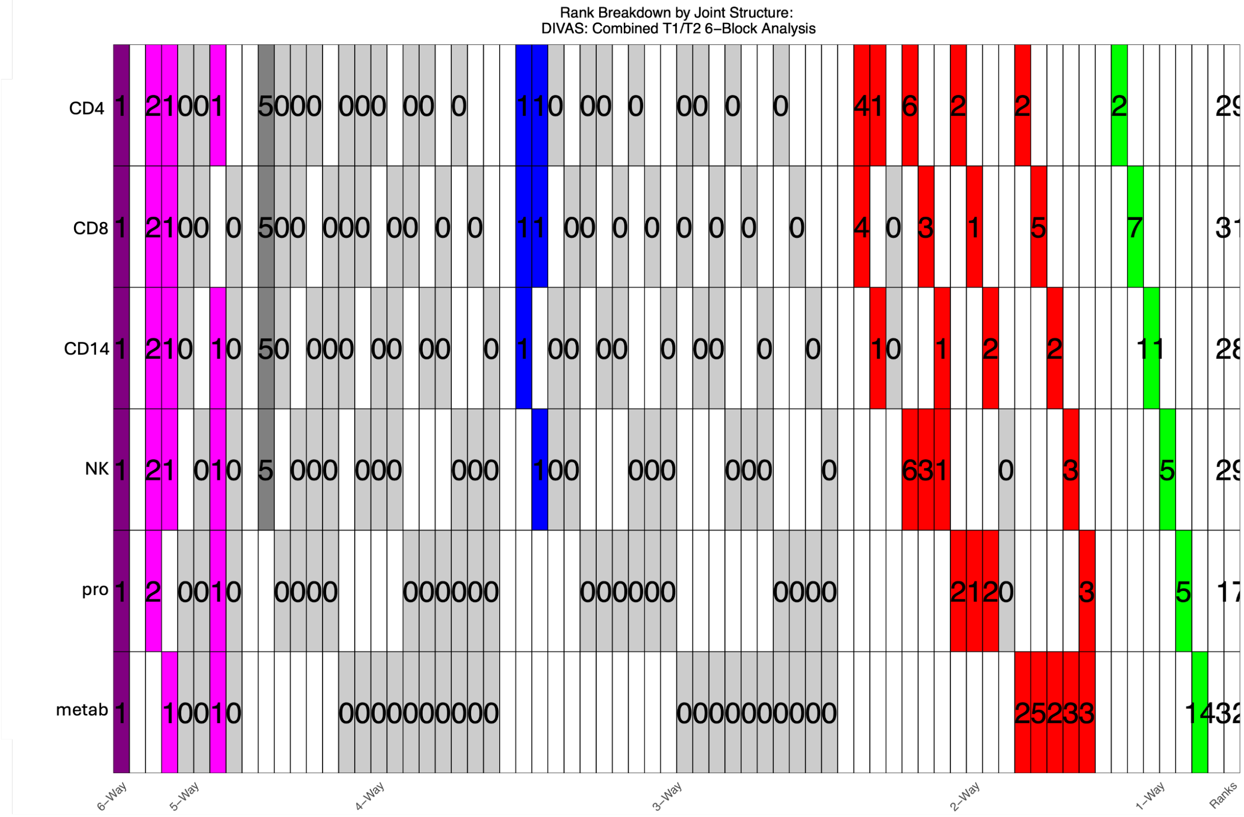


**Fig S1: Rank decomposition for the COVID-19 case study**. Each row represents one of the six modalities. The right most column prints the total number of components identified in each modality. All components were categorised into 6-way (purple), 5-way (pink), 4-way (dark gray), 3-way (blue), 2-way (red) and 1-way (green) variations. Each column represents a specific combination of modalities. For example, the first column (purple) represents the 6-way jointly shared components; the second column (pink) represents the CD4-CD8-CD14-NK-Prot 5-way partially shared components. In this case, one 6-way and two CD4-CD8-CD14-NK-Prot 5-way components were identified. This figure is produced directly by `DIVASmain`, which returns the rank decomposition among its diagnostic outputs when `iprint` is set or `figdir` is supplied; no separate plotting call is required.

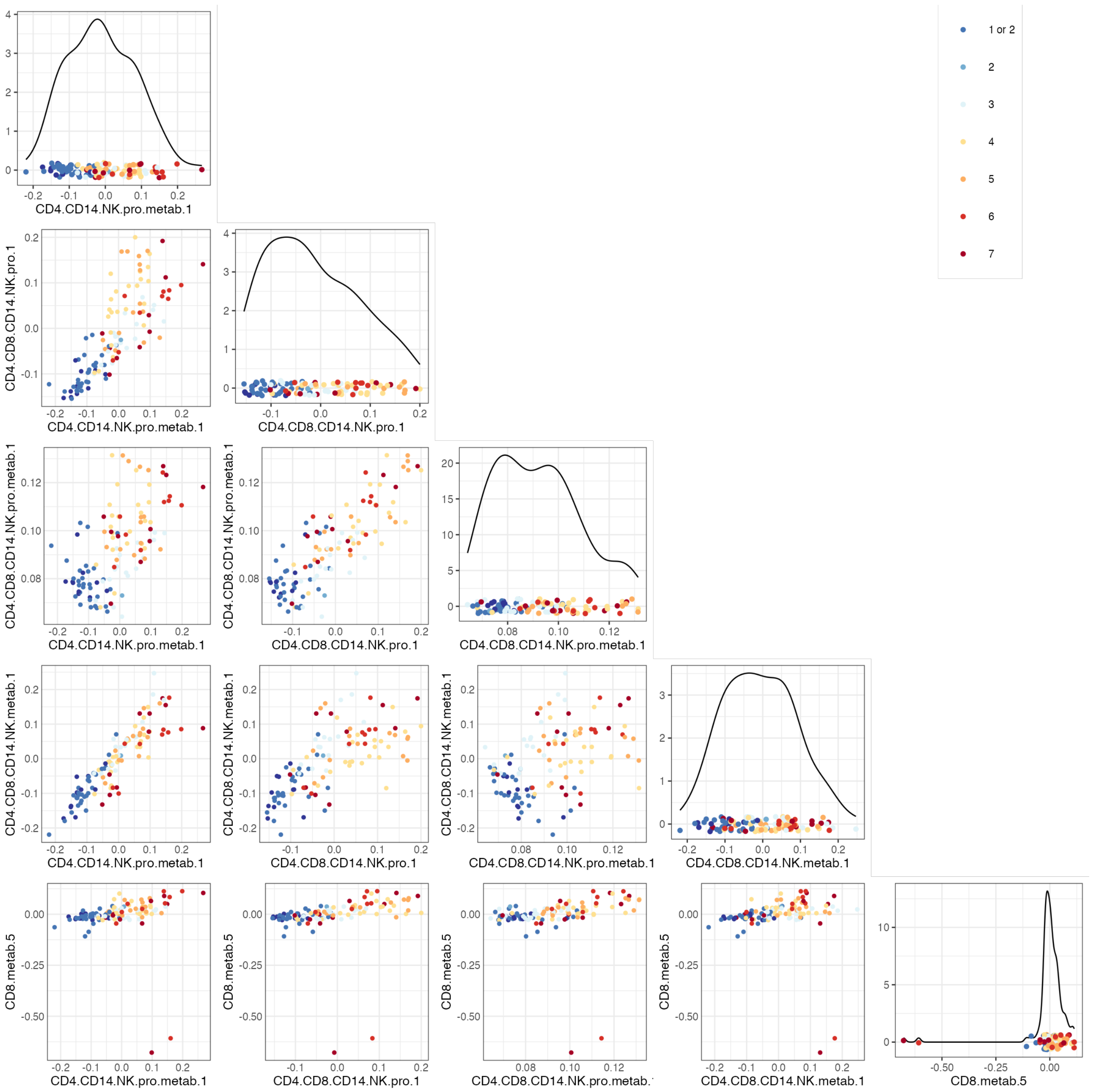


**Fig S2: Top 5 DIVAS components most strongly correlated with COVID-19 severity**. Probability density of the scores of the 5 components plotted on the diagonal. Other panels are scatter plots of pairwise combinations of scores, with data points coloured by COVID-19 severity score.

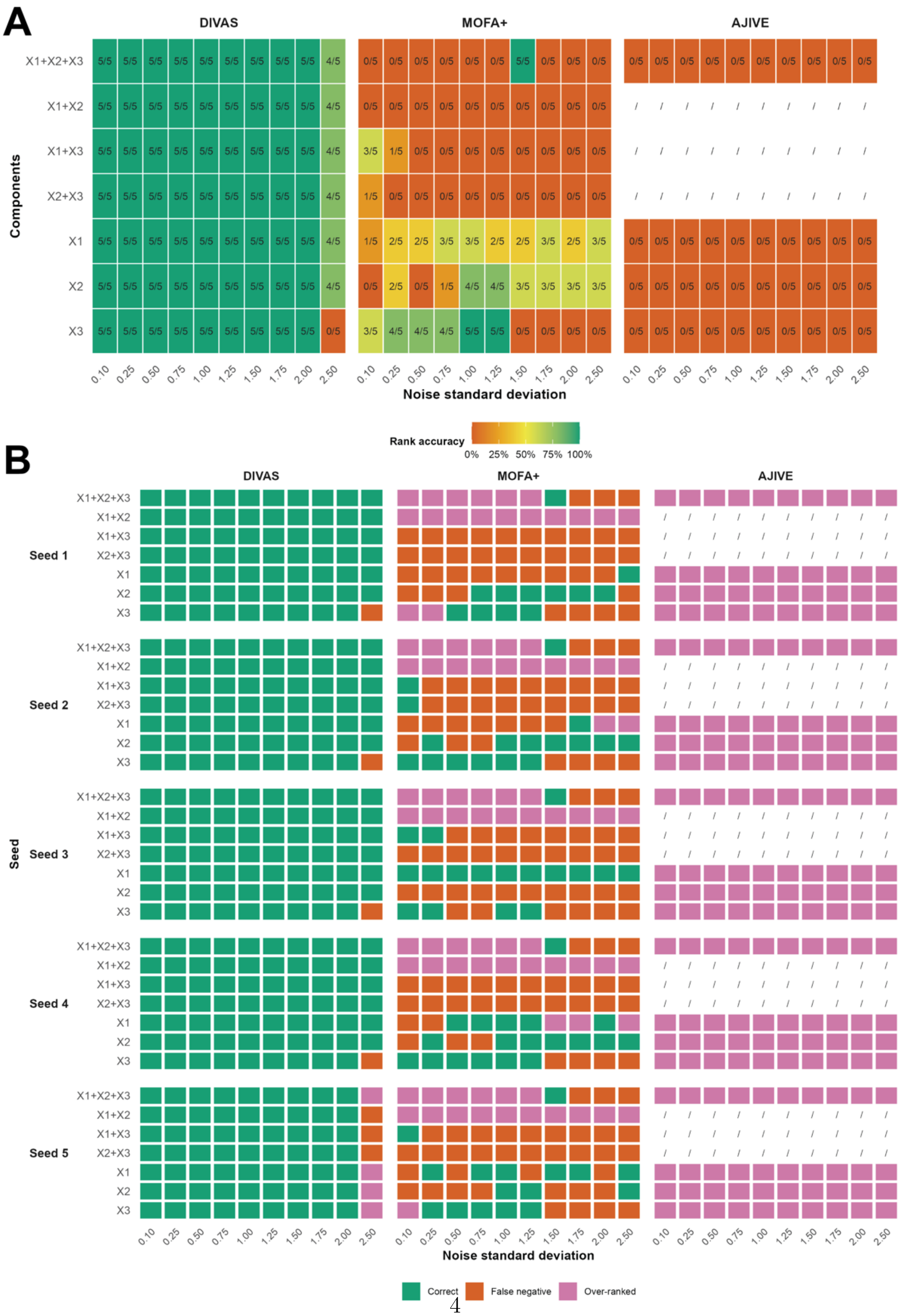


**Fig S3: Rank recovery in the generative benchmark for DIVAS, MOFA+ and AJIVE**. **A** Rank accuracy across ten noise standard deviations. Each tile reports the number of seeds, out of five, in which the rank of the corresponding index set was recovered exactly. **B** Rank recovery outcome for each of the five seeds, classified as correct, false negative (a true component with estimated rank zero) or over-ranked (an estimated rank larger than the true rank of one). In both panels, AJIVE is shown only for the jointly shared and individual components; the pairwise partially shared components are marked '/' because AJIVE does not define them. See Supplementary Methods Section S4.

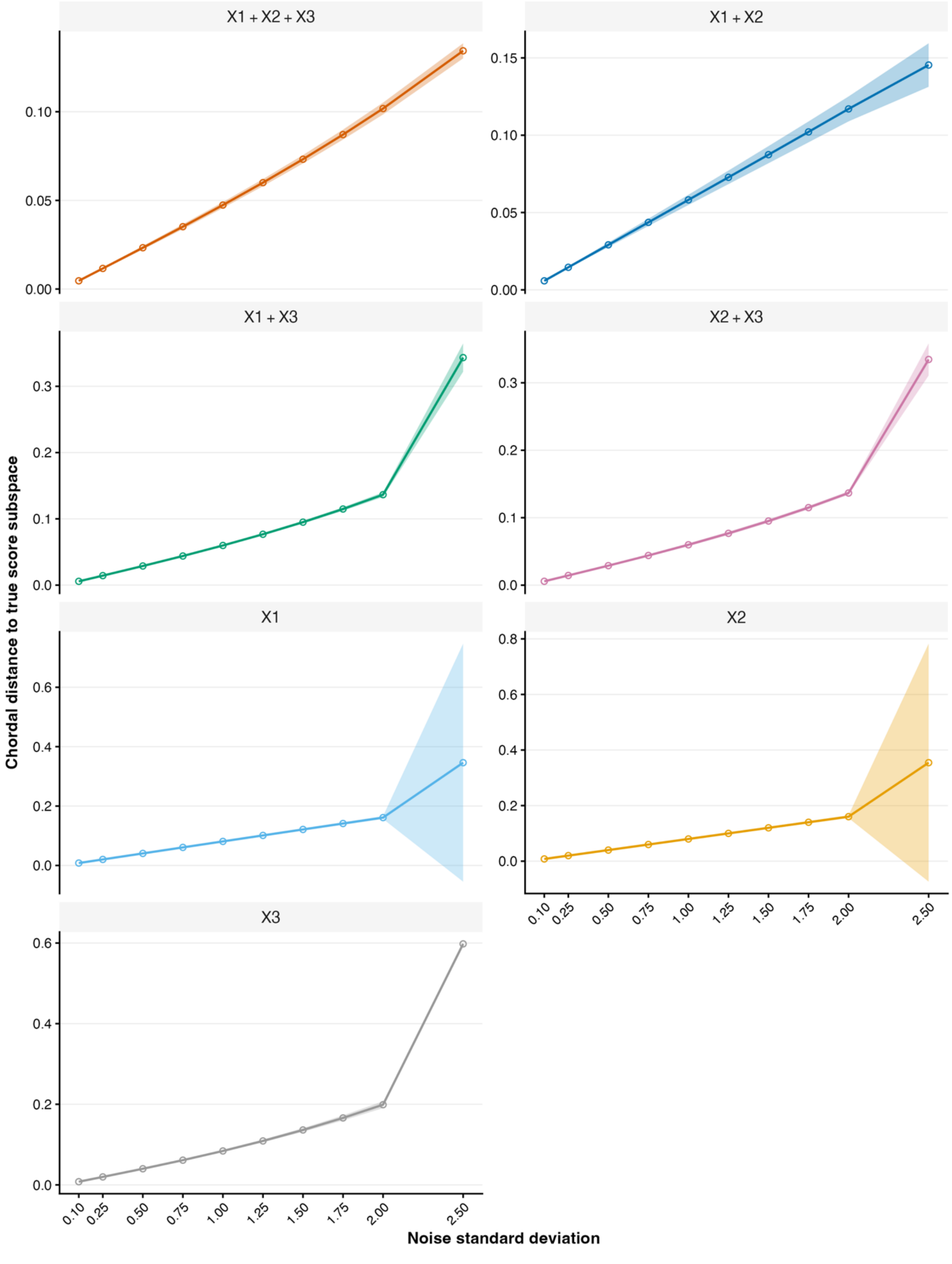


**Fig S4: DIVAS score-subspace recovery in the generative benchmark**. Chordal distance between the estimated and the true score subspace, for each of the seven active index sets, across ten noise standard deviations. Lines show the mean over five seeds and shaded bands the corresponding 95% confidence intervals. Smaller distances indicate closer recovery of the true generating score direction. See Supplementary Methods Section S4.

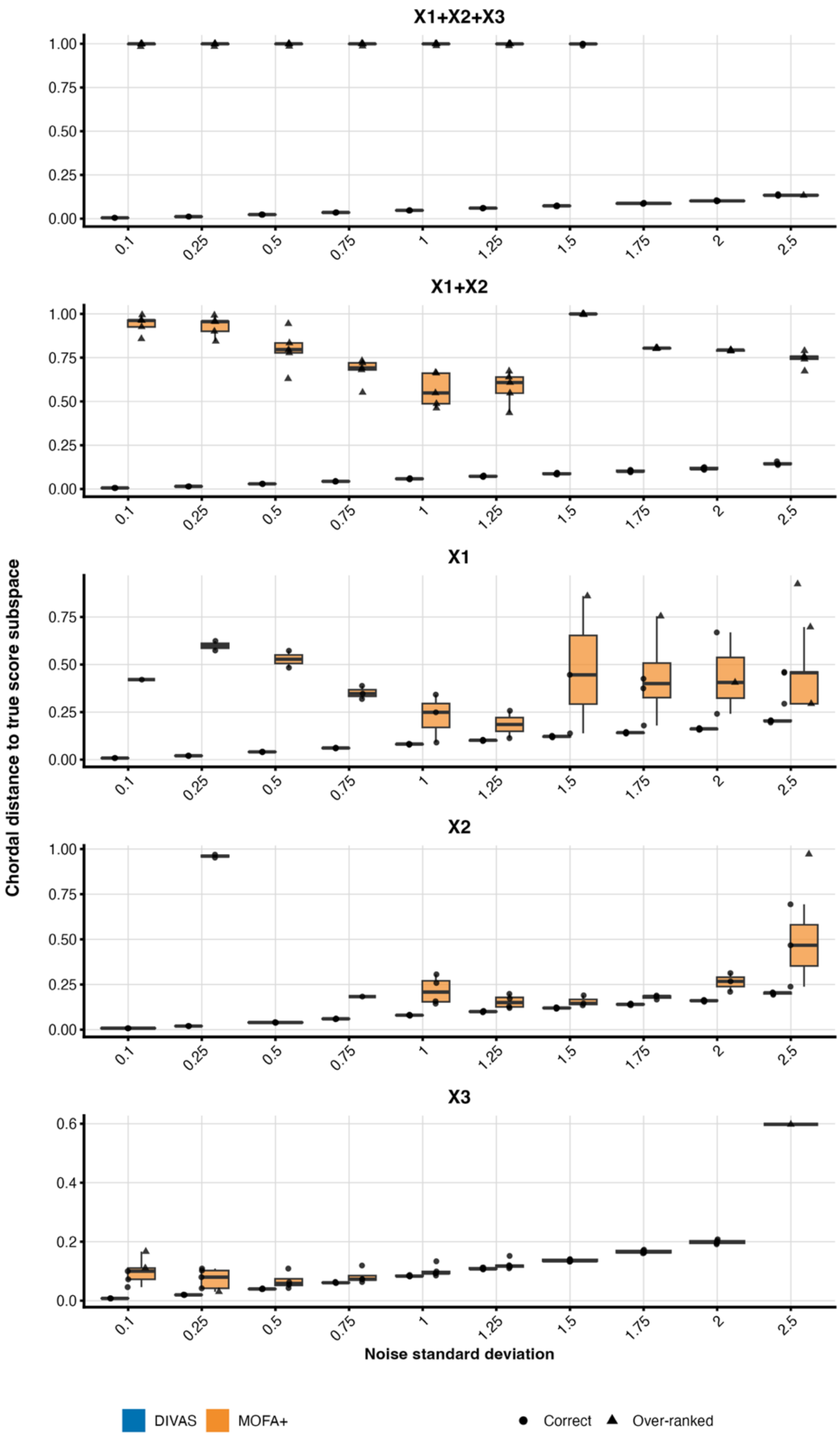


**Fig S5: Score-subspace recovery of DIVAS compared with mapped MOFA+ outputs in the generative benchmark**. Boxplots show the chordal distance to the true score subspace across five seeds and ten noise standard deviations. Only index sets with an evaluable mapped MOFA+ subspace are shown; $\{1, 2\}$ is the only pairwise partially shared subset for which MOFA+ produced sufficient mapped spans. Point shapes distinguish correct from over-ranked estimates. See Supplementary Methods Section S4.

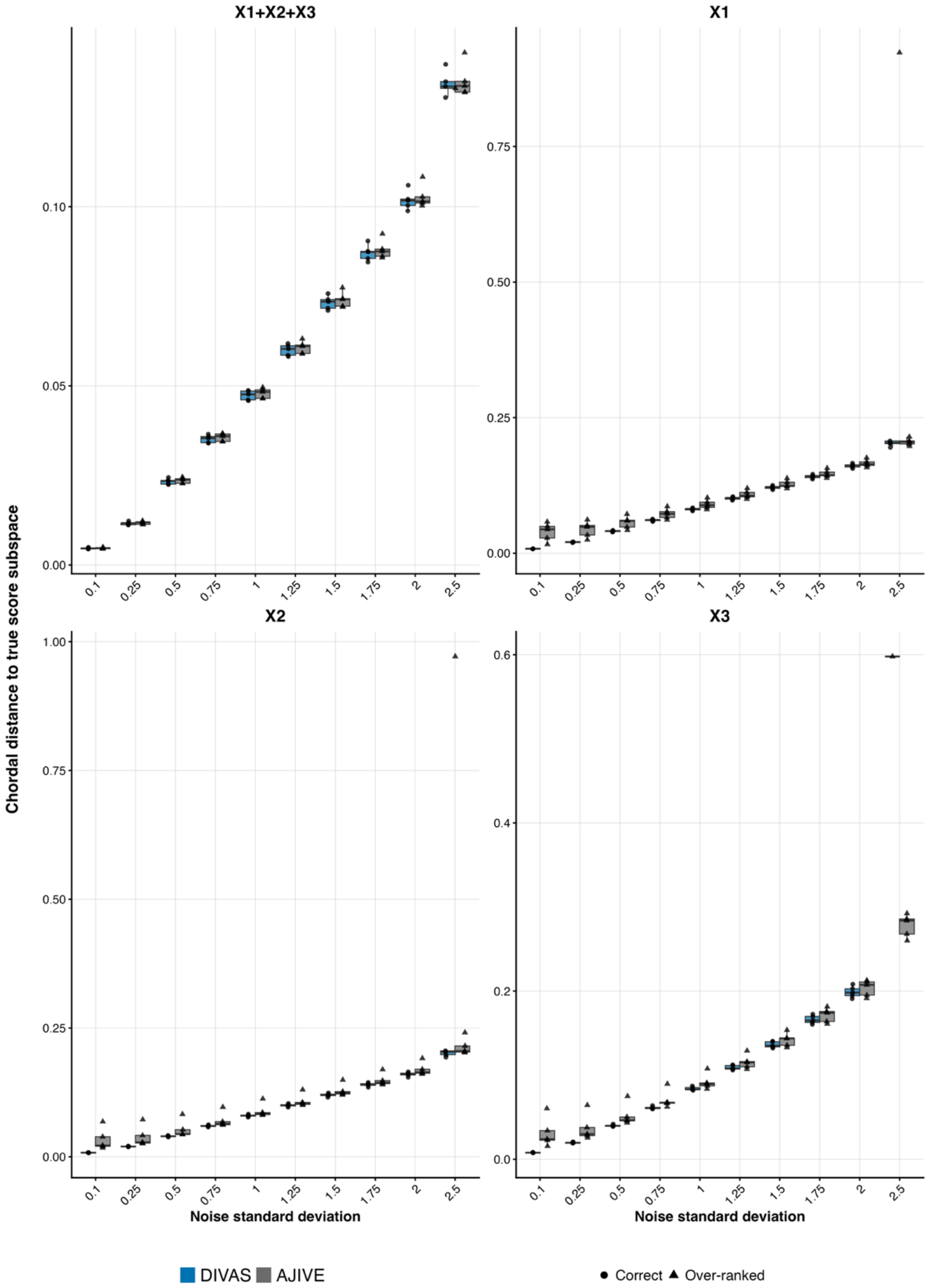


**Fig S6: Score-subspace recovery of DIVAS compared with AJIVE outputs in the generative benchmark**. Boxplots show the chordal distance to the true score subspace across five seeds and ten noise standard deviations. The pairwise partially shared subsets are omitted because AJIVE does not define them. Point shapes distinguish correct from over-ranked estimates; every AJIVE estimate was over-ranked. See Supplementary Methods Section S4.

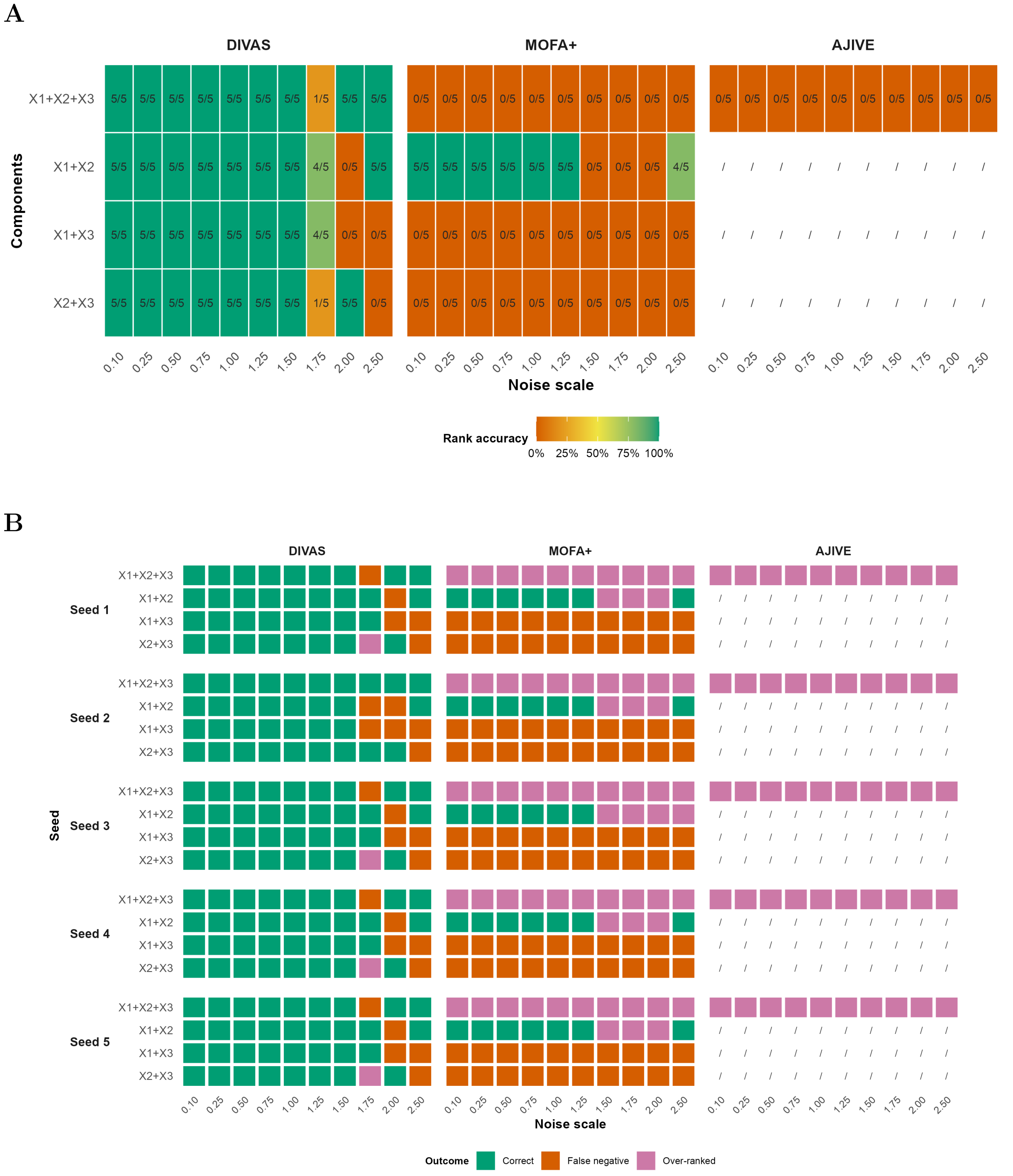


**Fig S7: Rank recovery in the additional toy benchmark for DIVAS, MOFA+ and AJIVE.** The published three-block `toyDataThreeWay` dataset contains one jointly shared and three pairwise partially shared components and no individual components. 'Noise scale' denotes the multiplier applied to the empirical standard deviation of each block. **A** Rank accuracy, with each tile reporting the number of seeds, out of five, with exact rank recovery. **B** Rank recovery outcome for each seed. Pairwise partially shared components are marked '/' for AJIVE, which does not define them. See Supplementary Methods Section S4.

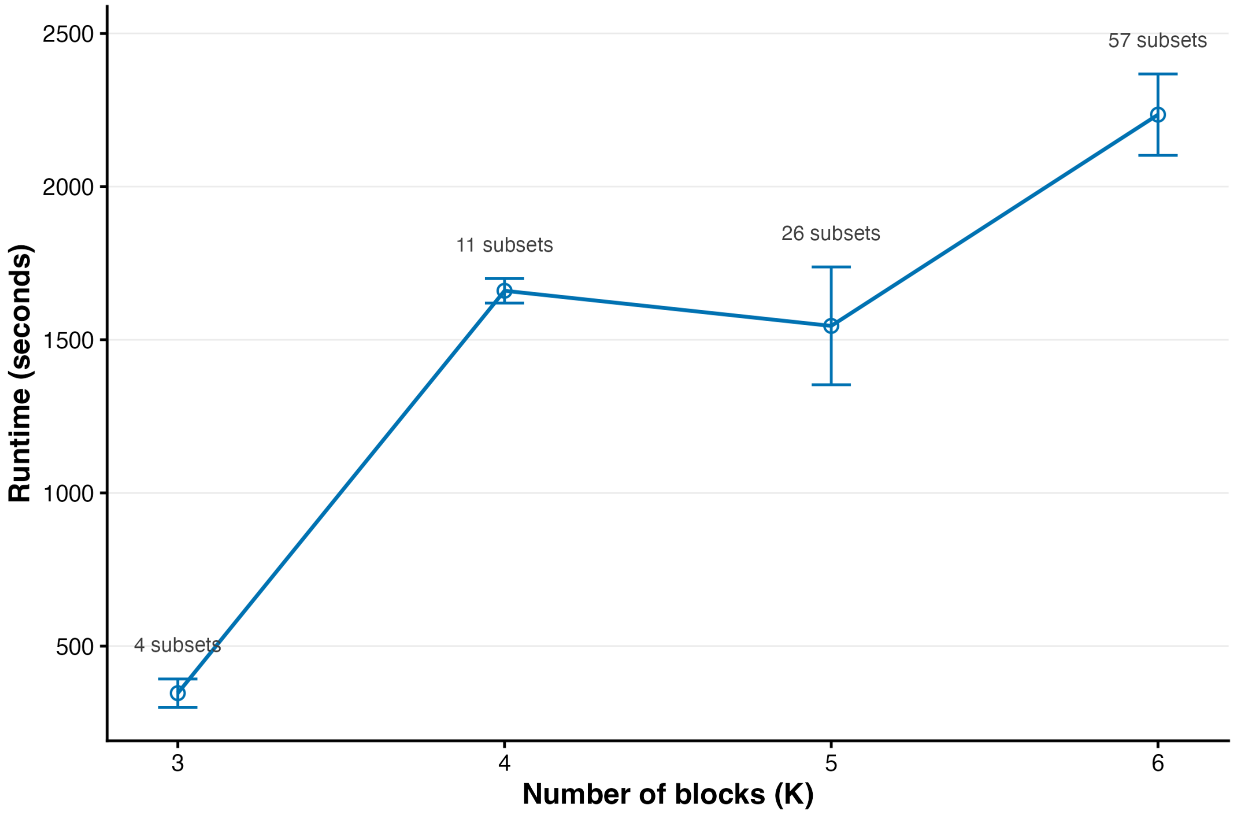


**Fig S8: DIVAS runtime as the number of data blocks increases**. Mean runtime over 10 seeds for simulations with $K = 3$ to 6 blocks; error bars show 95% confidence intervals. The annotation above each point gives the number of candidate sharing subsets searched by DIVAS at that $K$. All other simulation settings were held fixed; see Supplementary Methods Section S4.
