## Supplemental Methods for "DIVAS: an R package for identifying shared and individual variations of multiomics data"

#### S1 COVID-19 data preparation

**Data acquisition.** Raw multi-omics data were obtained from the COVID-19 study by Su et al. [1], which profiled 139 patients at two timepoints (T1: baseline; T2: follow-up), totally 265 patient samples. Single-cell RNA sequencing (scRNA-seq) data were downloaded from ArrayExpress under accession number E-MTAB-9357. Bulk proteomics and metabolomics data, along with clinical metadata, were obtained from the associated Mendeley Data repository (`https://data.mendeley.com/datasets/tzydswhhb5/5`). For the present DIVAS analysis, we focused on 120 patients with complete multi-omics measurements across both timepoints, resulting in 240 samples (120 patients times 2 timepoints).

**ScRNA-seq data preprocessing.** Raw scRNA-seq data were processed following quality control (QC) filtering: minimum 200 genes per cell, maximum 2,500 genes per cell, minimum 3 cells per gene and maximum 5 percent mitochondrial gene content. Cells passing QC were normalized to 10,000 counts per cell, followed by log1p transformation.

**Bulk proteomics and metabolomics preprocessing.** For metabolomics data, QC consisted of first removing metabolites with greater than 20 percent missing values across all samples. Remaining missing values in the retained metabolites were then imputed using the k-nearest neighbours (knn) algorithm with $k = 10$. Probabilistic quotient normalisation (PQN) was subsequently applied to correct for sample-to-sample variations in dilution. For proteomics data, an identical QC workflow was implemented. After QC, the final datasets comprised 481 proteins and 763 metabolites across 240 samples.

**Cell type annotation and pseudo-bulking.** Next, to enable cell type-specific analysis, we performed automated cell type annotation on the quality-filtered scRNA-seq data using CellTypist [2, 3] with the 'Adult COVID-19 PBMC' model and its build-in function of majority voting to assign cell type labels.

Following cell type annotation, we focused on four major immune cell populations relevant to COVID-19 pathogenesis: CD4 positive T cells (combining CD4 memory and naive subtypes), CD8 positive T cells (combining CD8 memory and naive subtypes), CD14 positive classical monocytes, and natural killer (NK) cells. For each cell type and each sample, we calculated the mean gene expression profile across all cells of that type, generating cell type-specific pseudo-bulk expression matrices. To ensure data quality, we applied further gene filtering: genes were retained only if they were expressed (non-zero) in at least 10 percent of samples within each time point (T1 or T2), and genes were further restricted to those present in both time points. This filtering resulted in 8,634 common genes across all four cell types.

**Sample selection to break autocorrelation.** From the 120 patients (240 samples) with complete multi-omics data, we retained exactly one observation per patient, so that no patient contributes repeated measurements. Stratified sampling was used to split the 120 patients into two groups of 60, balanced across key clinical variables: COVID-19 severity score (ranging from 1 to 7), disease severity categories (healthy, mild, moderate and severe), sex, age groups and ethnicity. The T1 observation was then retained for the first group and the T2 observation for the second, giving 120 candidate samples spanning both time points. Of these, 114 had complete measurements in all six data blocks and were retained for DIVAS analysis. This balanced design optimises statistical power for detecting associations between multi-omics patterns and clinical outcomes while controlling for potential confounding factors.

**Final DIVAS input data blocks.** We thus prepared 6 data blocks, each measured on the same 114 samples: 4 cell-type specific gene expression blocks (CD4 T, CD8 T, CD14 monocyte, NK; each with 8,634 genes) and two bulk omics blocks (proteomics with 481 proteins and metabolomics with 763 metabolites). All data blocks were aligned to ensure identical sample ordering across modalities. Row-wise centring (ensuring zero mean for each feature) was applied to each data block prior to DIVAS analysis.

All preprocessing scripts and intermediate data files are available in the GitHub repository (`https://github.com/ByronSyun/DIVAS_COVID19_CaseStudy`). Data files including the final DIVAS input matrices (Combined_CD4_T_combined.csv, Combined_CD8_T_combined.csv, Combined_CD14_Monocyte.csv, Combined_NK.csv, Combined_proteomics.csv, Combined_metabolomics.cs and pre-computed DIVAS results (divas_results_combined_6block_renamed.rds) are hosted on Zenodo (DOI: 10.5281/zenodo.17430294). Cell-level metadata for all 480,984 annotated cells (all_cells_metadata_complete.csv) is also available on Zenodo. Sample-level clinical metadata (metadata.rds) containing 31 clinical variables for the 114 samples is also included in the GitHub repository.

### S2 Downstream analyses

**Gene set overrepresentation.** To identify biological pathways and processes associated with selected DIVAS components in the gene expression (GEX) modalities (CD4 T, CD8 T, CD14 monocyte and NK cells), we performed gene set overrepresentation analysis using the `clusterProfiler` package (version 4.18.1) in R. For each component of interest and a GEX modality (e.g. CD4 GEX), we extracted the top 100 genes with the largest positive loadings and top 100 features with the largest negative loadings from each modality-specific loading vector. These two sets of features represent molecular signatures associated with opposing directions of the component scores.

We conducted gene ontology (GO) enrichment analysis using the `enrichGO` function from clusterProfiler. The analysis was performed against the biological process (BP) ontology using the human genome annotation database (`org.Hs.eg.db`). Gene symbols were used as the identifier type (`keyType = 'SYMBOL'`). Statistical significance was assessed using the default hypergeometric test with Benjamini-Hochberg correction for multiple testing. Enriched GO terms with adjusted p-value $< 0.05$ were considered statistically significant.

For proteomics and metabolomics modalities, where feature identifiers did not conform to standard gene nomenclature, we employed an alternative approach. Top-ranked features from these modalities were manually curated and their biological significance was interpreted through systematic literature review.

**Circos plot.** To visualise cross-modality feature relationships for selected DIVAS components, we generated circos plots using the `circlize` package (version 0.4.16) in R. We focused on the top-ranking component most strongly associated with COVID-19 severity (CD4–CD14–NK–pro–metab_1). From each of the five constituent modalities, we extracted the top 50 features with the largest positive loadings using `DIVAS::getTopFeatures`.

To identify cross-modality relationships, we calculated pairwise Spearman correlation coefficients between all features from different modalities using the preprocessed, row-centred data matrices. Correlations exceeding asymmetric thresholds were retained for visualisation: positive correlations $\geq 0.75$ and negative correlations $\leq -0.4$. These asymmetric thresholds were chosen to capture both strong positive coordinated variation and moderate negative relationships whilst maintaining visualisation clarity. The circos plot was then constructed using `circlize::chordDiagram`.

### S3 Simulation benchmarking

#### S1 Generative benchmark

To benchmark both rank recovery and score-subspace recovery under known ground truth, we generated a three-block simulated dataset with known score vectors, loading vectors and additive Gaussian noise, following the low-rank structure assumed by DIVAS (Section 2 of the main text).

We simulated $K = 3$ matched data blocks $X_1, X_2, X_3$ with $N = 400$ samples and $(d_1, d_2, d_3) = (200,\ 400,\ 10000)$ features. The underlying signal structure was specified by the collection of index sets

$$\mathcal{S}_{\text{true}} = \big\{\{1,2,3\},\ \{1,2\},\ \{1,3\},\ \{2,3\},\ \{1\},\ \{2\},\ \{3\}\big\},$$

that is, one jointly shared signal $\{1,2,3\}$, three partially shared signals $\{1,2\}$, $\{1,3\}$ and $\{2,3\}$, and three individual signals $\{1\}$, $\{2\}$ and $\{3\}$, each of true rank one. The $k$th block was generated as

$$X_k = \sum_{\boldsymbol{i} \in \mathcal{S}_{\text{true}}:\, k \in \boldsymbol{i}} \lambda\, l_{\boldsymbol{i},k} v_{\boldsymbol{i}}^{\top} + E_k, \qquad E_{k,ij} \overset{\text{i.i.d.}}{\sim} N(0, \sigma^2),$$

where $v_{\boldsymbol{i}} \in \mathbb{R}^{N \times 1}$ is the unit-norm score vector shared by all blocks in $\boldsymbol{i}$, $l_{\boldsymbol{i},k} \in \mathbb{R}^{d_k \times 1}$ is the unit-norm loading vector for block $k$ within $\boldsymbol{i}$, and $\lambda$ is the signal strength. We used $\lambda = 250$ and $\sigma \in \{0.10,\ 0.25,\ 0.50,\ 0.75,\ 1.00,\ 1.25,\ 1.50,\ 1.75,\ 2.00,\ 2.50\}$, and repeated each configuration across five random seeds $(1, \ldots, 5)$, giving 50 simulated datasets.

Recall from Section 2 of the main text that DIVAS estimates the signal matrix of the $k$th block as $A_k = \sum_{\boldsymbol{i}:\, k \in \boldsymbol{i}} L_{\boldsymbol{i},k} V_{\boldsymbol{i}}^{\top}$, where $V_{\boldsymbol{i}} \in \mathbb{R}^{N \times r_{\boldsymbol{i}}}$ is the orthonormal score matrix shared by all blocks in $\boldsymbol{i}$ and $L_{\boldsymbol{i},k} \in \mathbb{R}^{d_k \times r_{\boldsymbol{i}}}$ is the corresponding orthonormal loading matrix for block $k$. We write $V_{\boldsymbol{i}}$ and $L_{\boldsymbol{i},k}$ rather than $v_{\boldsymbol{i}}$ and $l_{\boldsymbol{i},k}$ because an estimated rank $r_{\boldsymbol{i}}$ may exceed the true rank of one.

#### S2 Baseline methods

As reviewed in the main text, no other off-the-shelf method recovers partially shared variations in multimodal data in the way DIVAS does. For benchmarking we selected AJIVE [4] and MOFA+ [5], run on the same simulated datasets, random seeds and noise grid. AJIVE distinguishes only jointly shared from individual variations and therefore provides an example of model misspecification, since it was not designed for partially shared variation; MOFA+ does not explicitly model partially shared variation, but its output can be post-processed to recover it. The purpose of this benchmark was therefore not to evaluate the native objectives of these baselines, but to ask whether their outputs can be repurposed to recover the same subset-labelled structure that DIVAS targets. Table S1 summarises the outputs evaluated for each method.

**Table S1:** Outputs evaluated for DIVAS, MOFA+ and AJIVE in the simulation benchmark.

| Method | Output evaluated | Comments |
|---|---|---|
| DIVAS | Subset-specific ranks and scores, returned directly | Directly targets partially shared structures |
| MOFA+ | Rank: number of latent factors mapped to each active-view pattern. Score: span of the sample-level factor values for those factors | Does not estimate subset ranks; subset structure inferred post hoc by factor-to-view mapping |
| AJIVE | Ranks and scores of the global joint and the individual subspaces, returned directly | Provides global joint and individual components, but no pairwise partially shared components |

AJIVE was run using the `ajive` R implementation [6]. AJIVE decomposes multiblock data into a structure shared by all blocks and block-specific individual structures, but it does not define pairwise partially shared components such as $\{1, 2\}$, $\{1, 3\}$ or $\{2, 3\}$. In our evaluation, the AJIVE global joint score subspace was compared with the jointly shared component $\{1, 2, 3\}$, and the pairwise partially shared subsets were recorded as not available with mapped rank zero. When AJIVE estimated a joint rank larger than the true rank, a small chordal distance was interpreted as containment of the true generating direction within a larger estimated span, and was reported together with the rank error.

MOFA+ was run using the `MOFA2` package [7]. MOFA+ estimates global latent factors with block-specific weights and reports factor activity through the variance explained in each block. Because MOFA+ does not natively return ranks for predefined sharing subsets, we mapped each factor to a subset using a transparent rule: for every factor (`MOFA2::get_factors`) we extracted the proportion of variance explained in each block (`MOFA2::get_variance_explained`), marked a block as active for that factor if the variance explained exceeded 1%, and assigned the factor to the index set of its active blocks.

Factors sharing the same set of active blocks were merged, and the mapped rank of a subset was defined as the number of factors assigned to it. The merged factor columns were used as the estimated score subspace for chordal distance comparison. These mapped ranks and subspaces are post hoc quantities introduced solely to enable comparison against the DIVAS-style ground truth, and should not be interpreted as native MOFA+ outputs.

#### S3 Evaluation metrics

For each simulated dataset, each method yields an estimated rank for the index set $\boldsymbol{i}$,

$$\hat{r}_{\boldsymbol{i}} = \dim\left(\widehat{V}_{\boldsymbol{i}}\right),$$

where $\widehat{V}_{\boldsymbol{i}}$ is the estimated score matrix. The ground-truth rank is $r_{\boldsymbol{i}} = 1$ for the seven index sets in $\mathcal{S}_{\text{true}}$ and $r_{\boldsymbol{i}} = 0$ otherwise. Rank recovery was evaluated for every non-empty $\boldsymbol{i} \subseteq \{1, 2, 3\}$ using

$$\text{rank error}(\boldsymbol{i}) = |\hat{r}_{\boldsymbol{i}} - r_{\boldsymbol{i}}|, \qquad \text{rank accuracy}(\boldsymbol{i}) = 1 - \min\left\{|\hat{r}_{\boldsymbol{i}} - r_{\boldsymbol{i}}|,\, 1\right\}.$$

We classified each outcome as *correct* ($\hat{r}_{\boldsymbol{i}} = r_{\boldsymbol{i}}$), a *false negative* ($r_{\boldsymbol{i}} > 0$ but $\hat{r}_{\boldsymbol{i}} = 0$), a *false positive* ($r_{\boldsymbol{i}} = 0$ but $\hat{r}_{\boldsymbol{i}} > 0$) or *over-ranked* ($\hat{r}_{\boldsymbol{i}} > r_{\boldsymbol{i}} > 0$).

For index sets with a recovered estimated subspace, score-subspace recovery was evaluated using the principal-angle-based chordal distance [8]. Writing $\theta_1, \ldots, \theta_{r_{\boldsymbol{i}}}$ for the principal angles between the true score subspace, spanned by the columns of $V_{\boldsymbol{i}}$, and the estimated score subspace, spanned by the columns of $\widehat{V}_{\boldsymbol{i}}$, the chordal distance is

$$d_c\big(V_{\boldsymbol{i}}, \widehat{V}_{\boldsymbol{i}}\big) = \sqrt{\sum_{i=1}^{r_{\boldsymbol{i}}} \sin^2 \theta_i}.$$

Principal angles were computed from the singular values of $Q_{\boldsymbol{i}}^{\top} \widehat{Q}_{\boldsymbol{i}}$, where $Q_{\boldsymbol{i}}$ and $\widehat{Q}_{\boldsymbol{i}}$ are orthonormal bases of the true and estimated score subspaces respectively. A smaller chordal distance indicates closer recovery of the true generating score direction. Because every active index set has true rank one in this benchmark, $d_c = \sin\theta_1$, and the distance measures whether the true score direction is contained in the estimated span. It is therefore interpreted alongside the rank error rather than as a substitute for rank recovery: a method that over-estimates the rank can achieve a small chordal distance simply because the true direction lies inside a larger estimated span.

#### S4 Simulation results

**Generative benchmark.** DIVAS recovered the correct rank for all seven active index sets, in all five seeds, for every noise level up to $\sigma = 2.00$ (Fig S3). At the most extreme noise level, $\sigma = 2.50$, DIVAS remained correct in four of five seeds for six of the seven index sets, but failed to recover the individual component $\{3\}$ in all five seeds. The corresponding score-subspace results (Fig S4) show a low chordal distance that increases smoothly with noise: at $\sigma = 2.00$ the mean chordal distance ranged from 0.10 for the jointly shared component $\{1, 2, 3\}$ to 0.20 for the individual component $\{3\}$. At $\sigma = 2.50$ the widened confidence intervals for $\{1\}$ and $\{2\}$ reflect increased variability, with four of five seeds retaining the correct rank and a low chordal distance while the remaining seed over-estimated the rank and showed substantially degraded subspace alignment.

MOFA+ showed limited and inconsistent recovery after active-view mapping. Among the pairwise partially shared subsets (Fig S3A), mapped rank recovery was sparse: $\{1, 3\}$ was correctly mapped only at the two lowest noise levels (4 of 50 datasets), $\{2, 3\}$ was recovered once (1 of 50), and $\{1, 2\}$ was never correctly ranked (0 of 50) and was consistently over-ranked. The jointly shared component $\{1, 2, 3\}$ was correctly ranked in only 5 of 50 datasets. Recovery of the individual components was better but still unstable, ranging from 23 to 25 of 50 datasets. Chordal distances were compared only for subsets with an evaluable mapped subspace, that is $\hat{r}_i > 0$ (Fig S5); $\{1, 2\}$ was the only pairwise subset with sufficient mapped spans for this comparison. Even for $\{1, 2\}$, the mapped MOFA+ factor spans were much further from the true score subspace than the DIVAS estimates (mean chordal distance 0.78 versus 0.07). For $\{1\}$ and $\{2\}$, MOFA+ distances were also larger and more variable than those of DIVAS; for $\{3\}$ the mean distances were occasionally comparable, but rank recovery and stability across seeds and noise levels remained inferior to DIVAS.

AJIVE showed a different limitation. Because it estimates a global joint subspace and block-specific individual subspaces only, the pairwise subsets $\{1, 2\}$, $\{1, 3\}$ and $\{2, 3\}$ are not applicable. For the subsets that AJIVE does return, $\{1, 2, 3\}$, $\{1\}$, $\{2\}$ and $\{3\}$, it consistently over-estimated the rank, at every noise level and in every seed (Fig S3). AJIVE chordal distances were sometimes close to those of DIVAS, particularly for the jointly shared component (Fig S6), but this does not amount to correct recovery of the rank-one structure: the small distance indicates only that the true score direction was contained within a larger estimated AJIVE span. Overall, AJIVE spans could contain some of the true jointly shared and individual directions, but AJIVE could not recover the pairwise partially shared structures and therefore did not recover the complete subset decomposition.

**Additional toy benchmark.** As a further robustness check, we evaluated rank recovery on the published three-block toy dataset `toyDataThreeWay` from the original DIVAS paper [9], after adding independent Gaussian noise with standard deviation equal to a multiple (the 'noise scale') of the empirical standard deviation of each block. This dataset contains one jointly shared and three pairwise partially shared components and no individual components. Because the released data do not include the true latent score vectors, this analysis was restricted to rank recovery and was not used for score-subspace recovery. The results (Fig S7) were broadly consistent with the generative benchmark: DIVAS recovered all four components in all five seeds up to a noise scale of 1.50 and degraded only at higher noise scales, MOFA+ recovered $\{1, 2\}$ at low noise but no other component, and AJIVE over-ranked the jointly shared component at every noise scale and returned no pairwise components.

**Scalability.** We additionally ran a scalability simulation for DIVAS to assess its practical behaviour as the number of blocks increases. Here $K = 3, 4, 5$ and 6 matched blocks were simulated with fixed block dimension $d_k = 200$, fixed sample size $N = 200$, noise standard deviation $\sigma = 0.50$, signal strength $\lambda = 180$, and $\texttt{nsim} = 400$, the number of bootstrap resamples used to infer the angle bounds (the package default). The active sharing design was simplified to one jointly shared component across all blocks plus two pairwise components, $\{1, 2\}$ and $\{K - 1, K\}$; no individual components were constructed. Under this design, the number of candidate sharing subsets searched by DIVAS increases from 4 for $K = 3$ to 11 for $K = 4$, 26 for $K = 5$ and 57 for $K = 6$.

Across 10 seeds, all runs completed and all true subsets were recovered with perfect rank accuracy for every $K$. Mean runtime increased from 346 s at $K = 3$ to 1660 s at $K = 4$, 1545 s at $K = 5$ and 2235 s at $K = 6$ (Fig S8). The non-monotone difference between $K = 4$ and $K = 5$ indicates that the realised runtime depends not only on $K$ but also on the optimisation path, the seed-specific data realisation and the exact subset dictionary explored in a given run. These absolute times should be interpreted only for this specific benchmark setting: changing the sample size, block dimension, signal strength or noise level would shift the runtime scale even at the same $K$. The benchmark is intended as practical guidance on computational cost as the number of blocks grows, rather than as a claim about universal runtime behaviour.

All code for reproducing the simulation benchmarks is available in the GitHub repository [10].

### References


[1] Yapeng Su, Daniel Chen, Dan Yuan, Christopher Lausted, Jongchan Choi, Chengzhen L Dai, Valentin Voillet, Venkata R Duvvuri, Kelsey Scherler, Pamela Troisch, Priyanka Baloni, Guangrong Qin, Brett Smith, Sergey A Kornilov, Clifford Rostomily, Alex Xu, Jing Li, Shen Dong, Alissa Rothchild, Jing Zhou, Kim Murray, Rick Edmark, Sunga Hong, John E Heath, John Earls, Rongyu Zhang, Jingyi Xie, Sarah Li, Ryan Roper, Lesley Jones, Yong Zhou, Lee Rowen, Rachel Liu, Sean Mackay, D Shane O'Mahony, Christopher R Dale, Julie A Wallick, Heather A Algren, Michael A Zager, ISB-Swedish COVID19 Biobanking Unit, Wei Wei, Nathan D Price, Sui Huang, Naeha Subramanian, Kai Wang, Andrew T Magis, Jenn J Hadlock, Leroy Hood, Alan Aderem, Jeffrey A Bluestone, Lewis L Lanier, Philip D Greenberg, Raphael Gottardo, Mark M Davis, Jason D Goldman, and James R Heath. Multi-omics resolves a sharp disease-state shift between mild and moderate COVID-19. *Cell*, 183(6):1479–1495.e20, 2020. URL `https://www.cell.com/cell/fulltext/S0092-8674(20)31444-6?_returnUR=`.

[2] C Domínguez Conde, C Xu, L B Jarvis, D B Rainbow, S B Wells, T Gomes, S K Howlett, O Suchanek, K Polanski, H W King, L Mamanova, N Huang, P A Szabo, L Richardson, L Bolt, E S Fasouli, K T Mahbubani, M Prete, L Tuck, N Richoz, Z K Tuong, L Campos, H S Mousa, E J Needham, S Pritchard, T Li, R Elmentaite, J Park, E Rahmani, D Chen, D K Menon, O A Bayraktar, L K James, K B Meyer, N Yosef, M R Clatworthy, P A Sims, D L Farber, K Saeb-Parsy, J L Jones, and S A Teichmann. Cross-tissue immune cell analysis reveals tissue-specific features in humans. *Science*, 376(6594):eabl5197, 2022.

[3] Chuan Xu, Martin Prete, Simone Webb, Laura Jardine, Benjamin J Stewart, Regina Hoo, Peng He, Kerstin B Meyer, and Sarah A Teichmann. Automatic cell-type harmonization and integration across human cell atlas datasets. *Cell*, 186(26):5876–5891.e20, 2023.

[4] Qing Feng, Meilei Jiang, Jan Hannig, and J S Marron. Angle-based joint and individual variation explained. *J. Multivar. Anal.*, 166:241–265, 2018.

[5] Ricard Argelaguet, Damien Arnol, Danila Bredikhin, Yonatan Deloro, Britta Velten, John C Marioni, and Oliver Stegle. MOFA+: a statistical framework for comprehensive integration of multi-modal single-cell data. *Genome Biol.*, 21(1):111, 2020.

[6] Iain Carmichael. *idc9/r.jive: First GitHub release*, 2020. URL `https://doi.org/10.5281/zenodo.4091755`. R package version 0.0.1.

[7] Ricard Argelaguet, Damien Arnol, Danila Bredikhin, and Britta Velten. *MOFA2: Multi-Omics*

*Factor Analysis v2*, 2025. URL https://www.bioconductor.org/packages/release/bioc/html/MOFA2.html. Bioconductor Release 3.22.

[8] John H Conway, Ronald H Hardin, and Neil J A Sloane. Packing lines, planes, etc.: packings in Grassmannian spaces. *Experimental Mathematics*, 5(2):139–159, 1996. URL https://doi.org/10.1080/10586458.1996.10504585.

[9] Jack Prothero, Meilei Jiang, Jan Hannig, Quoc Tran-Dinh, Andrew Ackerman, and J S Marron. Data integration via analysis of subspaces (DIVAS). *TEST*, 33:633–674, 2024. URL https://doi.org/10.1007/s11749-024-00923-z.

[10] Jiadong Mao and Yinuo Sun. *Finding multiomic markers and pathways underpinning COVID-19 severity using DIVAS*, 2025. URL https://byronsyun.github.io/DIVAS_COVID19_CaseStudy/. R Markdown Vignette.
